# Formation of long-term memory without short-term memory by CaMKII inhibition

**DOI:** 10.1101/2024.01.28.577644

**Authors:** Myung Eun Shin, Paula Parra-Bueno, Ryohei Yasuda

**Author notes:** Correspondence: RY.

## Abstract

Most of the current models of long-term memory consolidation require prior establishment of short-term memory. Here, we show that optogenetic or genetic inhibition of CaMKII, a kinase important for synaptic plasticity, in an inhibitory avoidance task impairs short-term memory at 1 h but not long-term memory at 1 d. Similarly, cortico-amygdala synaptic potentiation was more sensitive to CaMKII inhibition at 1 h but not at 1 d. These results strongly suggest that long-term memory does not require the prior formation of short-term memory and that CaMKII-dependent synaptic plasticity specifically regulates short-term memory, but not long-term memory, for avoidance memory.

## Introduction

It has been thought that the consolidation of long-term memory (LTM; memory lasting days) and remote memory (RM; permanent memory) requires the prior formation of short-term memory (STM; memory lasting hours) ^1^. Most of the current models of memory consolidation involve the initial establishment of STM by synaptic plasticity, followed by cellular consolidation over hours, and then time-dependent information transfer to different brain regions ^2–6^. However, STM and LTM have distinct sensitivity to pharmacological and genetic manipulations ^7–9^, suggesting they may be processed in parallel pathways. It is unknown to what degree LTM depends on STM, and whether these types of memory rely on distinct molecular mechanisms or forms of synaptic plasticity. Here, we examined the role of CaMKII, a kinase critical for long-term potentiation (LTP) of excitatory synapses and the formation of various forms of memory ^10–12^, on memory with different durations in an inhibitory avoidance task (IA) ^5,7,13–15^.

## Results

To optogenetically study the process of memory acquisition and consolidation in IA, we transduced excitatory neurons in the mouse amygdala with adeno associated virus (AAV) encoding light-inducible CaMKII inhibitor paAIP2 or its inactive mutant paAIP2neg (paAIP2[R5A/R6A]) under the *Camk2a* promoter, and implanted optical fibers ∼200-300 µm bilaterally above the lateral amygdala ^15^ (**Supplementary Fig. 1; Supplementary Table 1**). We used a custom IA apparatus that consists of bright and dark compartments separated by a door. When the door is open, an animal can freely move between the compartments while being tethered to a patch cable (**Fig. 1a**). For training, a mouse was placed in the bright compartment while the door was closed, and was allowed to explore the arena for 1 minute, after which the door opened. The mouse quickly crossed to the dark compartment almost always due to its natural preference. This was followed by door closing and the administration of a brief foot shock (0.5 mA for 2 s) ^12^. Following 2 training sessions (“2x training”, takes typically 6-8 min), the mouse was again placed in the bright compartment and its memory performance was quantified by measuring the cross latency to the dark compartment upon door opening. If the animal has learned, it should avoid crossing to the dark compartment, resulting in longer cross latency. CaMKII was inhibited by illuminating the amygdala through the implanted fibers during and after the training for 1 h ^12^. We measured STM (after the 1 h-laser illumination) and LTM (1 d after training), respectively.

**Figure 1.**
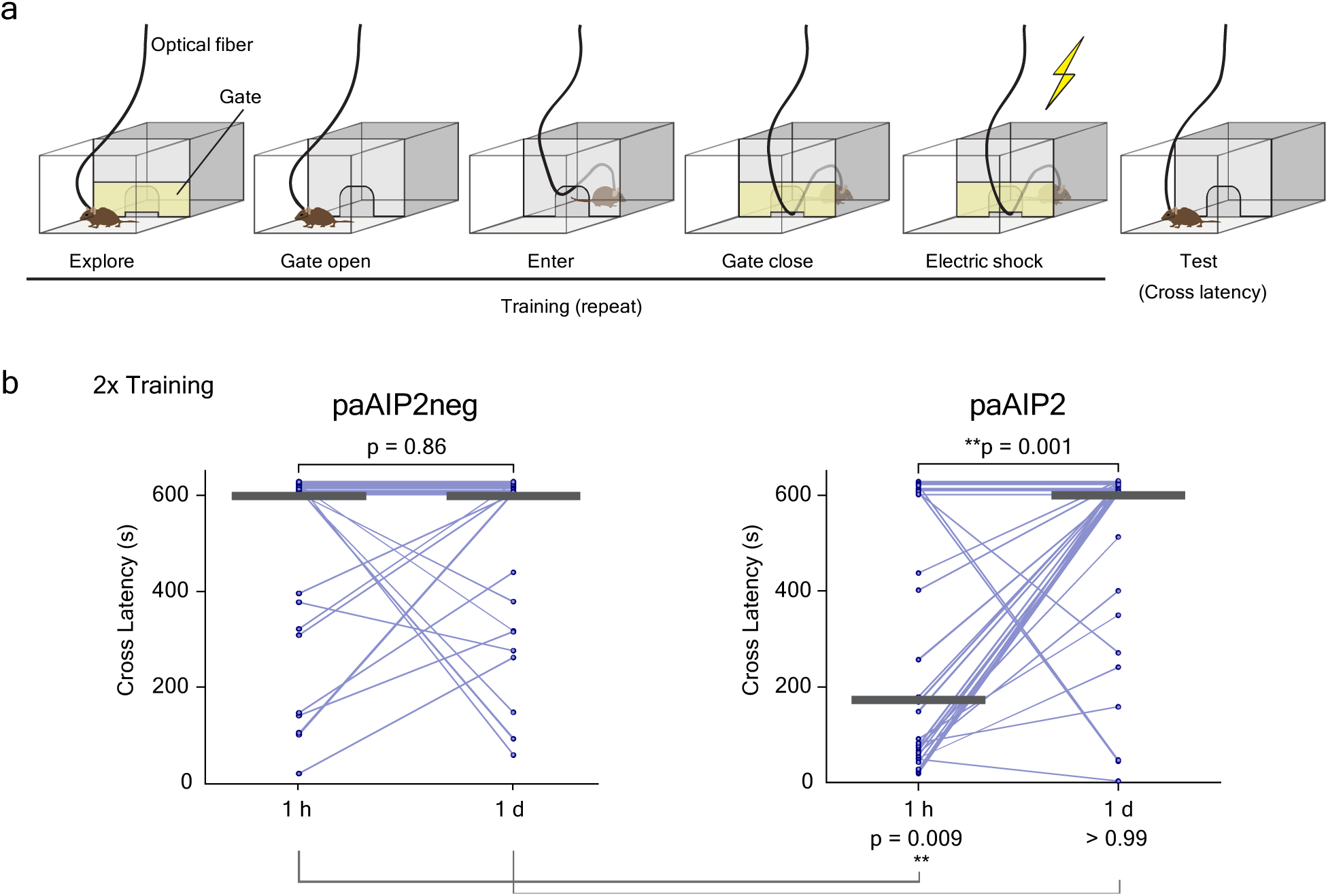
Performance of inhibitory avoidance task at different time points for mice in which CaMKII is transiently inhibited in amygdala. **a,** Schematics of inhibitory avoidance (IA) task. A mouse is allowed to explore the area for 1 minute, after which the door connecting the two chambers opens. When the animal crosses to the dark chamber, the door is closed and a foot shock is administered. The training is repeated once more. Each training trial ends when the animal crosses or avoids the dark chamber for 120 s. Mice were bilaterally injected with AAV encoding paAIP2neg (control) or paAIP2 into the amygdala, and implanted with optical fibers (800 μm Φ) ∼200-300 μm above the amygdala. Laser light was delivered to the amygdala during and after two trials of training for 1 h (473 nm, cycles of 1 s light on at 30 mW and 4 s off). **b,** Cross latency of IA test for mice injected with AAV encoding paAIP2neg (left) and paAIP2 (right) into the amygdala. Light was turned off before the 1 h test. Median is indicated by the horizontal bar. Two-sided Wilcoxon rank sum test was performed between the two time points within each group. Mann-Whitney test was conducted between paAIP2neg and paAIP2 groups at each time points. n = 27 for paAIP2neg and 33 for paAIP2 group. **p < 0.01 Data points with maximum latency values (600 s) have been randomized between 601 and 630 s to prevent overlapping.

For both paAIP2neg and paAIP2 groups, the cross latency increased between the first and second training, suggesting that memory acquisition is not affected by CaMKII inhibition in amygdala (**Supplementary Fig. 2**). The mice injected with paAIP2neg showed strong inhibition for crossing at both time-points, suggesting that their avoidance memory was intact in these animals (**Fig. 1b**). In comparison, the mice injected with paAIP2 showed strong impairment at 1 h. Remarkably, this impaired memory was recovered at 1 d in CaMKII-inhibited mice (**Fig. 1b**). The plot from individual animals in the paAIP2 group (n = 33) clearly shows that the majority of animals with a cross latency less than ∼300 s at the first testing trial (1 h) showed improvements on 1 d (18 out of 19 animals). These results suggest that CaMKII activation in the amygdala is required specifically for STM but not for LTM. Furthermore, the memory that was impaired at 1 h can be recovered in the later trials at 1 d.

Since multiple brain regions are likely involved in the formation of avoidance memory ^5,7^, we also used transgenic mice with a point mutation at a critical phosphorylation site of CaMKII (*Camk2a*^T286A^) ^11^. This point mutation strongly inhibits LTP and memory formation ^11,16,17^. Unlike the mice in which CaMKII is inhibited transiently only in the amygdala, the mutant mice with a chronic CaMKII inhibition in the entire brain showed impairment in memory acquisition during training (**Supplementary Fig. 3a**). However, by training 3-4 times, instead of 2 times, the mutant mice were able to learn the avoidance task (**Supplementary Fig. 3b, c**). For memory tests, we have measured immediate memory (IM) and remote memory (RM; 1 week and 4 weeks) as well as STM and LTM. When the mice received only 2x training, they failed to perform memory tests at all time points (**Supplementary Fig. 4**). However, they acquired a better overall memory after 3-4 training sessions (**Fig. 2**). In particular, after 4x training, the mice showed specific impairment only for 1 h memory (STM), but normal performance median levels for IM (2-3 min) and RM (7 d and 4 w).

**Figure 2.**
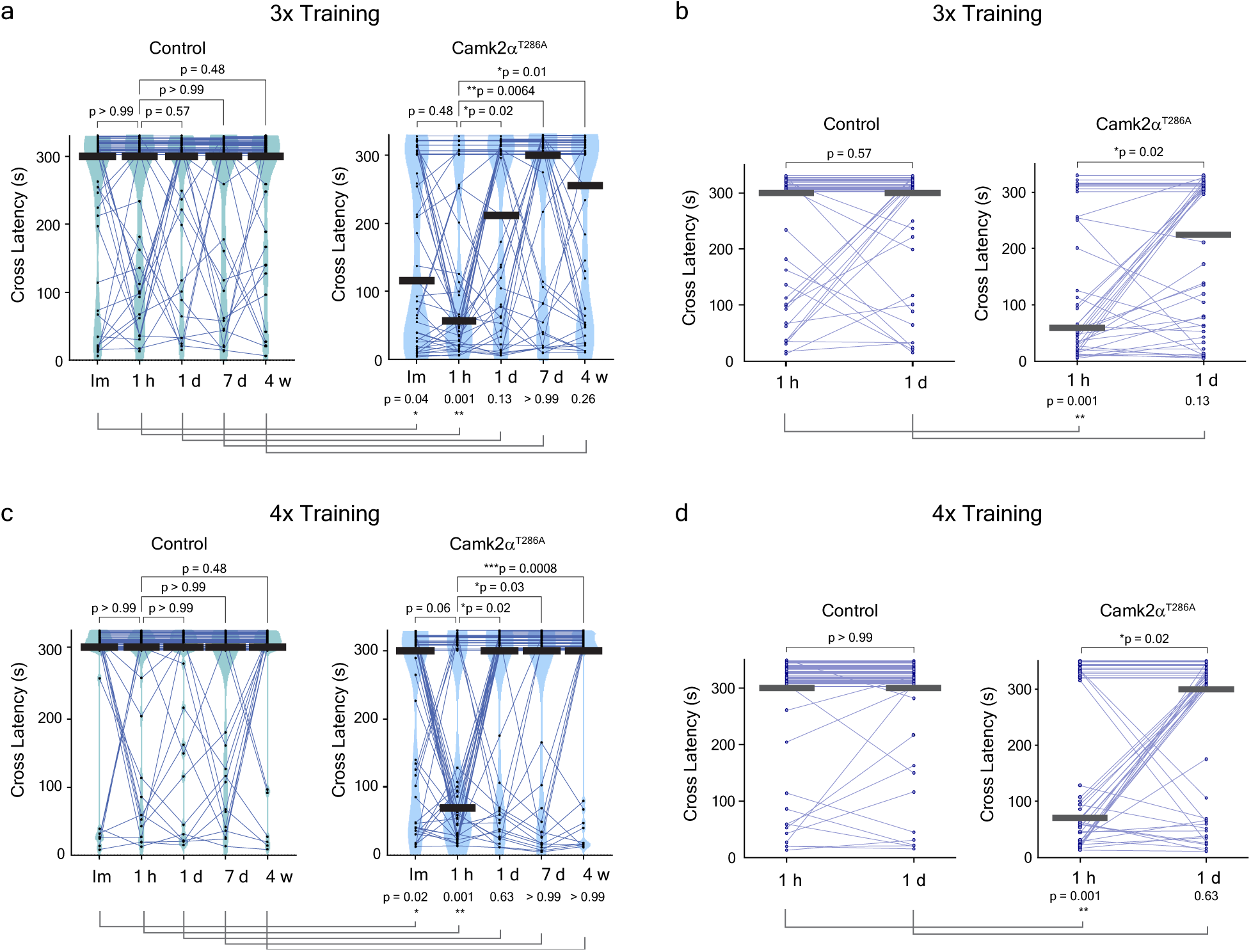
Performance of IA task for transgenic mice with inhibited CaMKII autophosphorylation. **a** and **c**, Performance of IA task immediately, 1 h, 1 d, 7 d and 4 w after 3 trials (**a**) or 4 trials (**c**) of training for *Camk2a*^T286A^ mice (right) or their littermate non-transgenic control (left). The experiment is similar to that in Fig. 1a, but without optical fibers. **b** and **d**, Cross latencies at two selected time points (1 h and 1 d) from **a** or **c** are shown, respectively. Median is indicated by the horizontal bar. Two-sided Wilcoxon rank sum test was performed between 1 h and the other respective time point(s) in each group. Mann-Whitney test was conducted between control and mutant mice at each time points. n = 40 for control and 41 for *Camk2a*^T286A^ mice (**a-b**), n = 47 for control and 45 for mutant mice (**c-d**). *p < 0.05, **p < 0.01, ***p < 0.001 Data points with maximum latency values (300 s) have been randomized between 301 and 330 s (**a-c**) or 350 s (**d**).

We performed two control experiments to confirm that the testing session did not cause memory recovery at 1 d. First, we tested if the time in the chamber for the memory test affected memory performance. We obtained similar results for 300 s and 600 s-maximum cross latency tests (**Fig. 2, Supplementary Fig. 5**). Next, to see if the memory test at 1 h influences the test result at 1 d, we skipped the 1 h test in a group of animals. There was no difference for the memory test at 1 d and 7 d between groups in which the 1 h test was performed or omitted (**Supplementary Fig. 6**). Overall, our results suggest that CaMKII-dependent plasticity is required to form STM but not for LTM or RM.

To find physiological correlates of the avoidance memory, we measured excitatory postsynaptic currents (EPSCs) in neurons of the lateral amygdala (LA), evoked by stimulating the cortico-amygdala fiber in *Camk2a*^T286A^ mutant mice and their wild-type (WT) littermate controls (**Fig. 3**). For 2X training, we saw a significant increase in AMPA receptor (AMPAR)- and NMDA receptor (NMDAR)-mediated current 1 h and 1 d after training in WT mice, consistent with behavioral change in avoidance. For the CaMKII mutant, however, we observed an increase in AMPAR current only at 1 d after the training but not at 1 h (**Fig. 3b**). For the intensive training (4x training), where the CaMKII mutant animals had impaired memory at 1 h but intact memory at 1 d, the cortico-amygdala synapses showed a significant potentiation in AMPAR current both at 1 h and 1 d (**Fig. 3d**). The AMPAR to NMDAR ratio did not exhibit appreciable differences in all the conditions we measured (**Fig. 3b, d; Supplementary Fig. 7**). Overall, the electrophysiological recording showed 1 h potentiation is more susceptible to CaMKII inhibition than 1 d potentiation, similar to the general trend of avoidance memory. However, electrophysiological readout of synaptic plasticity in response to avoidance training appears to be more sensitive than behavioral output since mutant animals that received 2x training show impaired avoidance but normal synaptic potentiation at 1 d after training.

**Figure 3.**
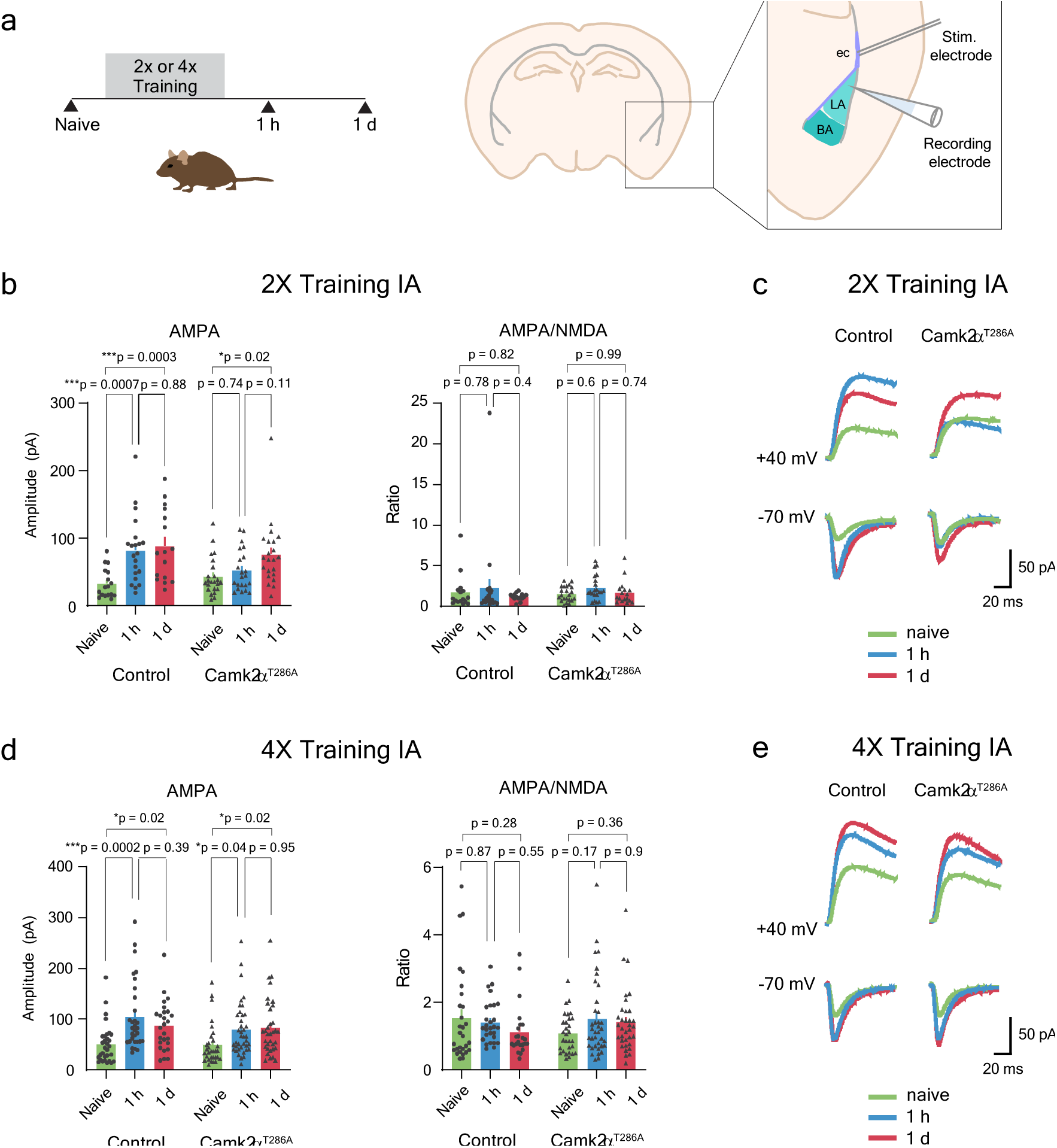
The effect of IA on the amplitude of AMPA and NMDA currents in acute brain slices obtained from transgenic mice with inhibited CaMKII autophosphorylation. **a**, Left: Experimental design. Coronal acute slices of brains were obtained from *Camk2a*^T286A^ mice and their littermate control that received 2 or 4 trials of training. Brains were collected 1 h and 1 d after training or from naïve mice with no prior training. Right: Whole cell recordings were performed in cells in lateral amygdala (LA) in acute slices. A stimulation electrode was placed in the external capsule (ec) to stimulate the cortical input to the LA. **b** and **d,** Whole cell recording after 2 trial- or 4 trial-training IA, respectively. Left: The amplitude of peak AMPA currents (calculated at -70 mV). Right: AMPA/NMDA ratios. Black triangles and black circles indicate individual data points for transgenic and control mice, respectively. Bars indicate mean with SEM. Two-way analysis of variance (ANOVA) was conducted. In **b**, for AMPA, interaction effect: F_2, 115_ = 2.63, p = 0.08; time effect F_2, 115_ = 12.3, p < 10^-4^; genotype effect F_1, 115_ = 2.13, p = 0.15. For AMPA/NMDA ratio, interaction effect: F_2, 106_ = 0.16, p = 0.85; time effect F_2, 106_ = 1.2, p = 0.31; genotype effect F_1, 106_ = 0.07, p = 0.8. In **d**, for AMPA, interaction effect: F_2, 186_ = 1.09, p = 0.34; time effect F_2, 186_ = 12.3, p < 10^-4^; genotype effect F_1, 186_ = 1.64, p = 0.2. For AMPA/NMDA ratio, interaction effect: F_2, 175_ = 2.32, p = 0.1; time effect F_2, 175_ = 0.66, p = 0.52; genotype effect F_1, 175_ = 0.02, p = 0.89. n_cells/animals_ for 2x IA AMPA = 17/6 (control naïve), 21/5 (control 1 h), 16/5 (control 1 d), 23/11 (mutant naïve), 22/8 (mutant 1 h), 22/8 (mutant 1 d). n_cells/animals_ for 2x IA AMPA/NMDA ratio = 16/6 (control naïve), 21/5 (control 1 h), 15/5 (control 1 d), 22/11 (mutant naïve), 21/8 (mutant 1 h), 17/6 (mutant 1 d). n_cells/animals_ for 4x IA AMPA = 31/7 (control naïve), 31/9 (control 1 h), 26/8 (control 1 d), 31/10 (mutant naïve), 39/11 (mutant 1 h), 34/13 (mutant 1 d). n_cells/animals_ for 4x IA AMPA/NMDA ratio = 29/6 (control naïve), 28/8 (control 1 h), 23/7 (control 1 d), 30/9 (mutant naïve), 37/11 (mutant 1 h), 34/13 (mutant 1 d). Tukey’s multiple comparisons; *p < 0.05, **p < 0.01, ***p < 0.001, ****p < 0.0001 **c** and **e**, Representative curves of mean AMPA and NMDA responses (at -70 mV and +40 mV, respectively) for control (left) and mutant (right) mice for 2 trial- and 4 trial-training IA, respectively.

## Discussion

This study revealed that transient CaMKII inhibition in the amygdala during training inhibits inhibitory avoidance memory at 1 h after the training but not at 1 d. For almost all mice that showed impaired memory at 1 h, we observed strong memory recovery between 1 h and 1 d. Furthermore, we also observed similar phenotypes in the mutant mice, in which CaMKII-dependent plasticity is chronically impaired in all Camk2a-positive neurons in the whole brain. Thus, our results suggest that CaMKII-dependent plasticity plays a specific role in the formation of STM but not in LTM for avoidance memory. Furthermore, CaMKII mutant animals showed a normal level of RM performance, suggesting that CaMKII inhibition does not affect memory maintenance and recall. The electrophysiological recording suggests that 2x training IA induces potentiation of cortico-amygdala connection, and this strengthening is impaired at 1 h but not at 1 d by CaMKII mutation, consistent with behavioral results. Since synaptic potentiation occured in LA in CaMKIIα mutant animals, even under a condition where behavioral output is impaired, there appears to exist CaMKIIα-independent plasticity that supports behavioral and synaptic plasticity. It has been proposed that LTM requires off-line LTP during sleep in the neocortex, while STM requires on-line LTP in the limbic system ^5^. Given the dissociation of STM and LTM phenotypes by CaMKII inhibition, off-line LTP may be CaMKII-independent and occur in parallel with on-line LTP.

## Methods

### Animals

All experimental protocols were approved and carried out in accordance with the regulations set out by Max Planck Florida Institute IACUC. Mice were housed under a reversed 12 h light/dark cycle with food and water *ad libitum*. Mice were between 2–5 months old and weighed approximately 27–35 g at the time of the behavior experiments. For optogenetic experiments, male mice (C57BL/6, Charles River) were randomly assigned to experimental (injected with paAIP2) and control (injected with paAIP2neg) groups. For experiments using transgenic mice (*Camk2a*^T286A^, gift from K. Peter-Giese, C57BL/6 background), homozygous mice and their littermate non-transgenic mice (both sexes) were used as experimental and control groups.

### Virus injection

AAV9 particles for CaMKIIp-mEGFP-P2A-paAIP2 (∼10^13^ infections units/ml) and its non-functional mutant CaMKIIp-mEGFP-P2A-paAIP2[R5AR6A] (paAIP2neg; ∼10^13^ infections units/ml) were packaged, purified and concentrated by the University of North Carolina Vector Core. Virus (1 μl/site) was bilaterally injected into the lateral amygdala (LA; From bregma, anteroposterior [AP] -1.65 mm, mediolateral [ML] ±3.75 mm, dorsoventral [DV] -3.8-3.9 mm) using a Picospritzer III (Parker) at a rate of 0.2 µl/min.

### Optical fiber implantation

Optical fiber implantation was performed with custom-made chronic implantable optical fibers (800 μm core Φ, 0.39 NA) housed inside a ferrule (2.5 mm OD, Precision Fiber Products). The fiber tips were placed ∼200-300 μm above BLA complex (AP -1.65 mm, ML ±3.75 mm, DV -3.6 mm) and the implant was secured to the skull using dental cement (Lang Dental). Optical fiber implantation was performed 2-3 weeks after viral injection. Behavioral experiments were conducted 4–6 weeks from viral injection. To activate paAIP2, blue light (0.2 Hz, 1 s, 30 mW output/site) from a laser (473 nm Blue DPSS Laser, Shanghai Laser & Optics Century) was delivered bilaterally through a custom-made bifurcating optical patch cable (200 μm core Φ, 0.22 NA) connected to the implanted optical fibers.

### Inhibitory avoidance task

The inhibitory avoidance (IA) task was performed following a previous published protocol ^15^. Briefly, we used a custom-made IA apparatus with two separated compartments with one room bright (500 lux) and the other dark. The apparatus allows free movement of an animal tethered to a patch cable. For optogenetic experiments, mice were handled while being attached with a dummy optical patch cable for 3 days (30–60 min/day) before the IA training. For experiments using transgenic mice, mice were handled for 3 days (5–10 min/day) prior to the IA training. The training was performed in the following steps. A mouse was placed inside the bright chamber, then allowed to explore and adapt to the light chamber for 1 min, after which the door opened. Once it crossed to the dark chamber, the door was closed. The “crossing” was defined as the simultaneous detection of three body points (nose-point, center point and tail-base) of the mouse beyond a small entry area bordering light chamber within dark chamber. The cross latency was defined as the time from door opening to crossing. After a short delay (6 s for the paAIP2 mice and 3 s for the transgenic mice), a mild foot shock (0.5 mA for 2 s) was administered to the mouse. 10 s after the foot shock, the training trial was finished. After a brief delay (∼2 min) for cleaning the apparatus, the training trial was repeated (total 2–4 times). If the mouse crossed to the dark chamber again within 120 s (cut-off point), the door was closed and the mouse received another foot shock. If the mouse stayed in the light chamber for 120 s, the training was finished. Mice were returned to the home cage once the training was completed. Subsequently, IA memory was assessed at various time points as follows. After ∼2 min of delay for cleaning the apparatus, immediate memory (IM) was tested by measuring the cross latency to the dark chamber. Short-term memory (STM) was assessed at 1 h from the training (transgenic mice) or after the 1 h-laser delivery (optogenetic mice). Long-term (LTM) and remote memories (RM) were assessed at twenty-four hours and 7 days and 4 weeks from the training, respectively. For testing sessions, 300 or 600 s cut-off point was applied as indicated in each figure. No electric shock or laser was applied during test sessions. Experimenter (MES) was blinded from genotypes of the mice or types of injected virus during behavioral experiments. Mice were excluded from the analysis if they did not cross for 120 s in the first training trial. Behavioral data was recorded via an infrared camera interfaced with Ethovision software (Noldus Information Technologies). After the experiments, infected areas in BLA were analyzed from mEGFP fluorescence in histology images. Analysis included animals expressing the mEGFP in BLA at least in one hemisphere.

### Histology, immunocytochemistry and microscopy

Mice were perfused transcardially with phosphate-buffered saline (PBS), followed by 4% paraformaldehyde (PFA) in PBS. Extracted brains were post-fixed in 4% PFA overnight. 70 μm coronal sections were obtained with a vibratome, washed in PBS three times for 10 min, stained for DAPI and mounted. Brain images were acquired on a Zeiss LSM 780 confocal microscope with a 10x objective (z stack, tiled). Obtained images were used to assess the expression of transgene and the location of the optical fiber track.

### Electrophysiology

Slices from *Camk2*α^T286A^ mice or their littermate WT control (2-3 months old) that received the 2x or 4x IA training were obtained at two different time points, 1 h and 1 d after training. Slices from naïve animals that received no training were collected as well. Mice were sedated by isoflurane inhalation and perfused intracardially with ice-cold Choline Chloride solution (Choline Chloride 124 mM; KCl 2.5 mM; NaHCO_3_ 26 mM; MgCl_2_ 3.3 mM; NaH_2_PO_4_ 1.2 mM; Glucose 10 mM; CaCl_2_ 0.5 mM; pH 7.4, equilibrated with 95% O_2_/ 5% CO_2_). Brains were extracted and placed in the same chilled solution for slicing. Then, 300 µm coronal slices containing the lateral amygdala were collected and placed in oxygenated (95% O_2_/ 5% CO_2_) ACSF (NaCl 127 mM; KCl 2.5 mM; Glucose 10 mM; NaHCO_3_ 25 mM; NaH_2_PO_4_ 1.25 mM; MgCl_2_ 2 mM; CaCl_2_ 2 mM) at 32°C for 1 h and maintained at room temperature for the whole cell recording experiment. Neurons in LA were recorded using whole cell voltage clamp using a Cs-based internal solution (CsMeth 120 mM; NaCl 10 mM; EGTA 0.3 mM; HEPES 10 mM; Mg ATP 4 mM; Na GTP 0.3 mM; QX314 5 mM; TEA-Cl 10 mM) and pipettes of 4-6 MΩ. To stimulate the incoming fibers for LA, a concentric bipolar stimulation electrode (FHC) was placed in the external capsule. We used a fixed stimulation strength (9V), which generated an EPSC of ∼50% of the maximal values for naïve animals. Neurons were clamped at -70 mV and then at +40 mV to determine the amplitude of the AMPA and NMDA current responses. The peak amplitude of the EPSC at -70 mV was calculated to determine the AMPA response, and the current measured 100 ms after the peak at +40 mV was used to determine the NMDA response.

### Statistical analysis

GraphPad Prism 10 software was used for statistical analysis. For inhibitory avoidance data, a two-sided Wilcoxon rank sum test was performed for assessing repeated measurements within a group and Mann-Whitney test was conducted for comparing unpaired measurements between two groups. Resulting p values were adjusted with Bonferroni’s correction. Two-way analysis of variance (ANOVA) was conducted for electrophysiological data. Statistical parameters such as the types of statistical tests, sample numbers (n) and statistical significance are reported in the figures and figure legends.

---

**Supplementary Figure 1.**
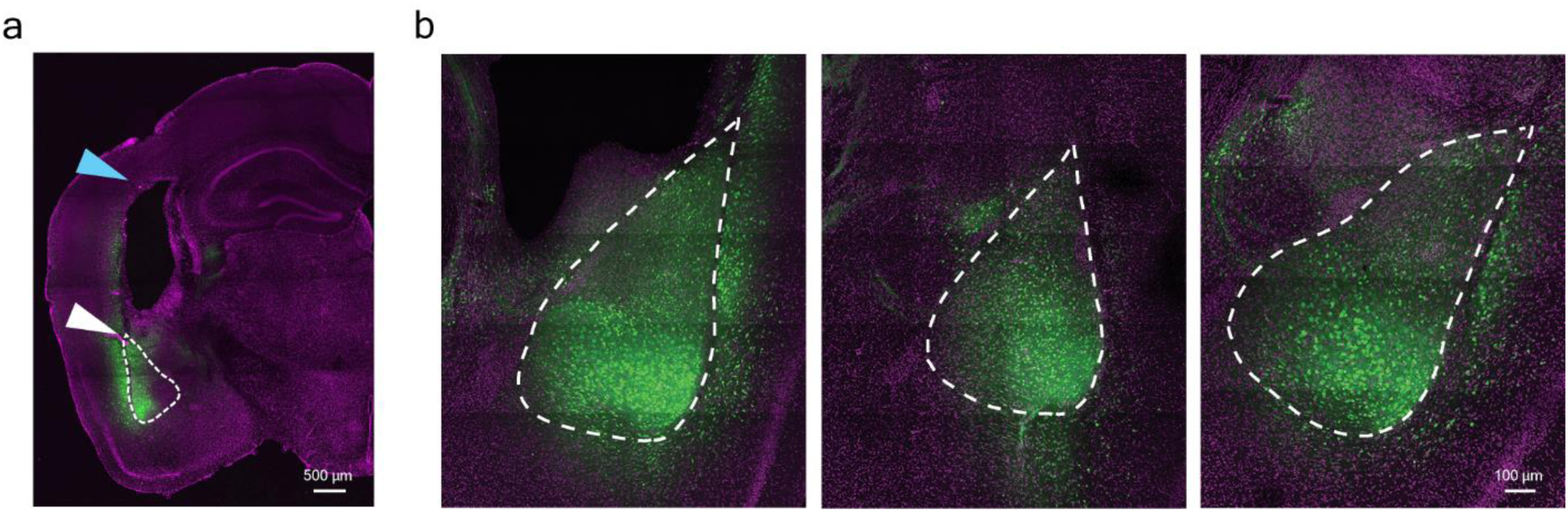
Expression of EGFP-P2A-paAIP2 in vivo. **a**, The expression of EGFP-P2A-paAIP2 in amygdala. The white and blue triangles indicate the area of transgene expression and the fiber track above amygdala, respectively. White dotted line indicates basolateral amygdala (BLA). Scale bar = 500 μm. Animals expressing detectable transgene expression in BLA in at least one hemisphere were included in the behavioural analyses. **b**, Examples of EGFP-P2A-paAIP2 expression in vivo following AAV injection. White dotted line indicates BLA. Scale bar = 100 μm. In some animals, the transgene expression was also found in the brain regions adjacent to the BLA, such as entorhinal cortex, piriform cortex, endopiriform nucleus and central and medial amygdala.

**Supplementary Figure 2.**
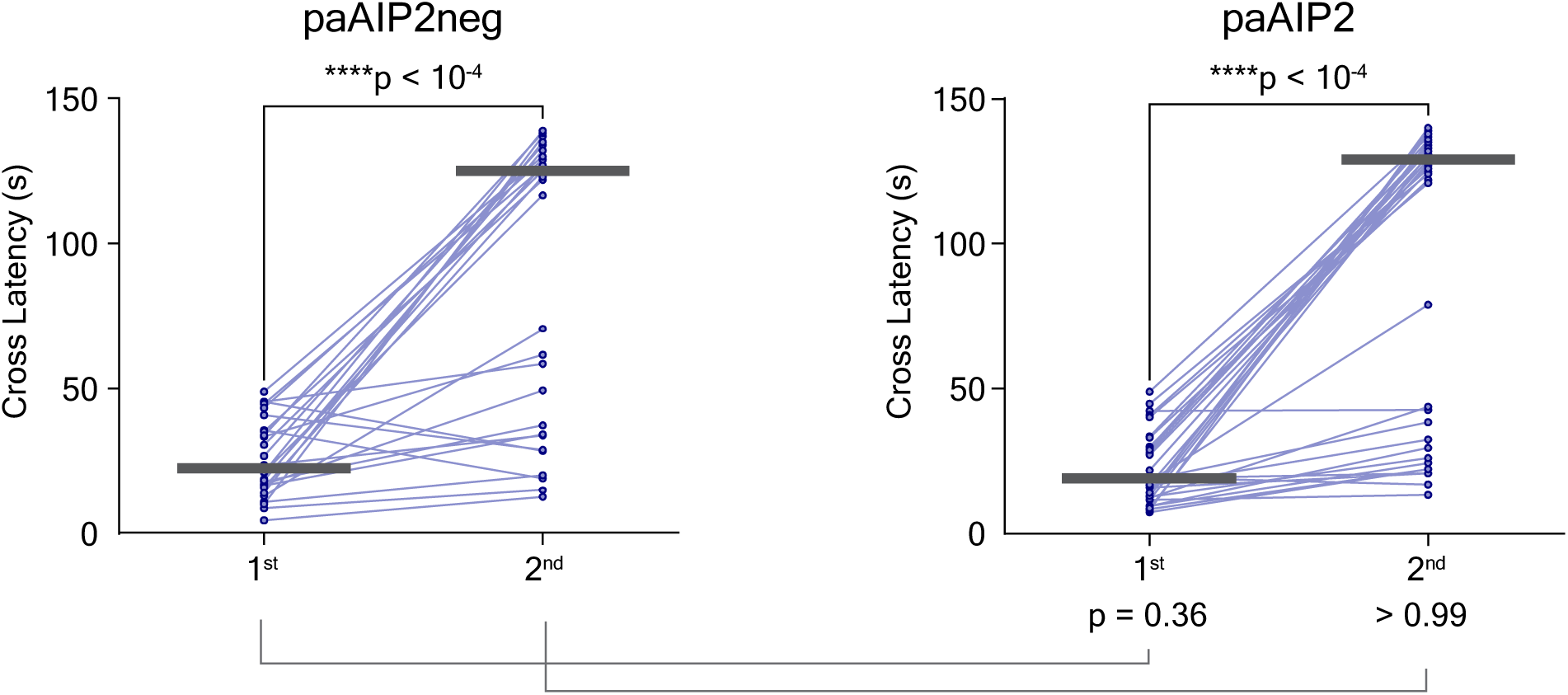
Acquisition of IA memory for mice in which CaMKII is transiently inhibited in amygdala. Cross latency during IA training for mice injected with paAIP2neg (left) or paAIP2 (right). Median is indicated by the horizontal bar. Two-sided Wilcoxon rank sum test was performed between the 2 trials of training in each group. Mann-Whitney test was conducted between paAIP2neg and paAIP2 groups at each trials. n = 27 for paAIP2neg and 33 for paAIP2 group. ****p < 0.0001 Data points with maximum latency values (120 s) have been randomized between 121 and 140 s.

**Supplementary Figure 3.**
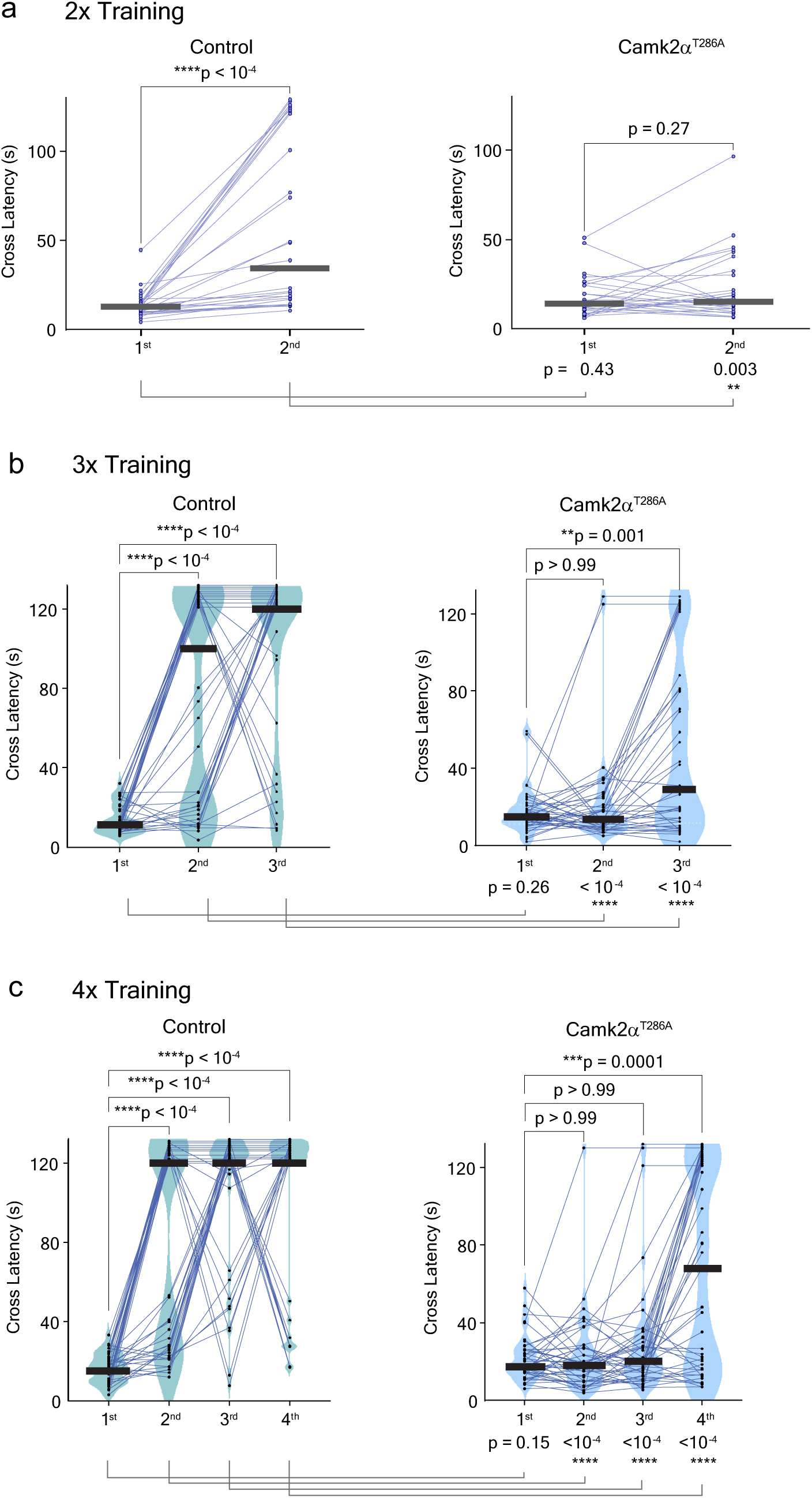
Acquisition of IA memory for transgenic mice with inhibited CaMKII autophosphorylation. **a-c**, Cross latency during multi-trial IA training for *Camk2a*^T286A^ mice (right) or their littermate non-transgenic control (left). Median is indicated by the horizontal bar. Two-sided Wilcoxon rank sum test was performed between the first and subsequent trials of training in each group. Mann-Whitney test was conducted between control and *Camk2a*^T286A^ mice at each trials. n = 29 for control and 28 for *Camk2a*^T286A^ mice (**a**), n = 40 for control and 41 for *Camk2a*^T286A^ mice (**b**), n = 47 for control and 45 for *Camk2a*^T286A^ mice (**c**). **p < 0.01, ***p < 0.001, ****p < 0.0001 Data points with maximum latency values (120 s) have been randomized between 121 and 132 s.

**Supplementary Figure 4.**
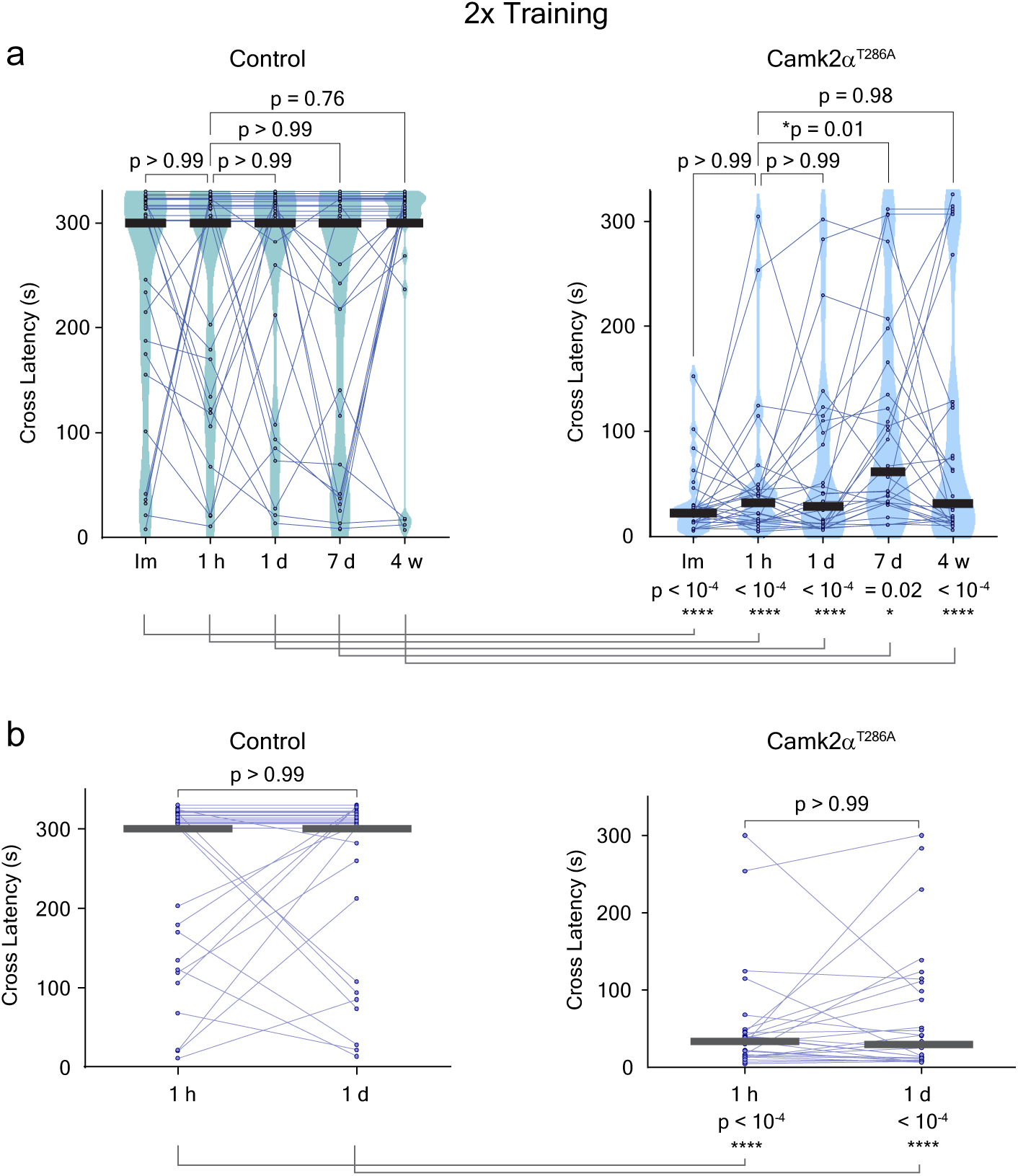
Performance of IA task with 2 trials of training for transgenic mice with inhibited CaMKII autophosphorylation. **a**, Performance of IA task immediately, 1 h, 1 d, 7 d and 4 w after 2 trials of training for *Camk2a*^T286A^ mice (right) or their littermate non-transgenic control (left). **b**, Cross latencies at two selected time points (1 h and 1 d) from **a** are shown. Median is indicated by the horizontal bar. Two-sided Wilcoxon rank sum test was performed between 1 h and the other respective time points in each group. Mann-Whitney test was conducted between control and transgenic mice at each time points. n = 29 for control and 28 for *Camk2a*^T286A^ mice (**a-b**). *p < 0.05, ****p < 0.0001 Data points with maximum latency values (300 s) have been randomized between 301 and 330 s (**a-b**).

**Supplementary Figure 5.**
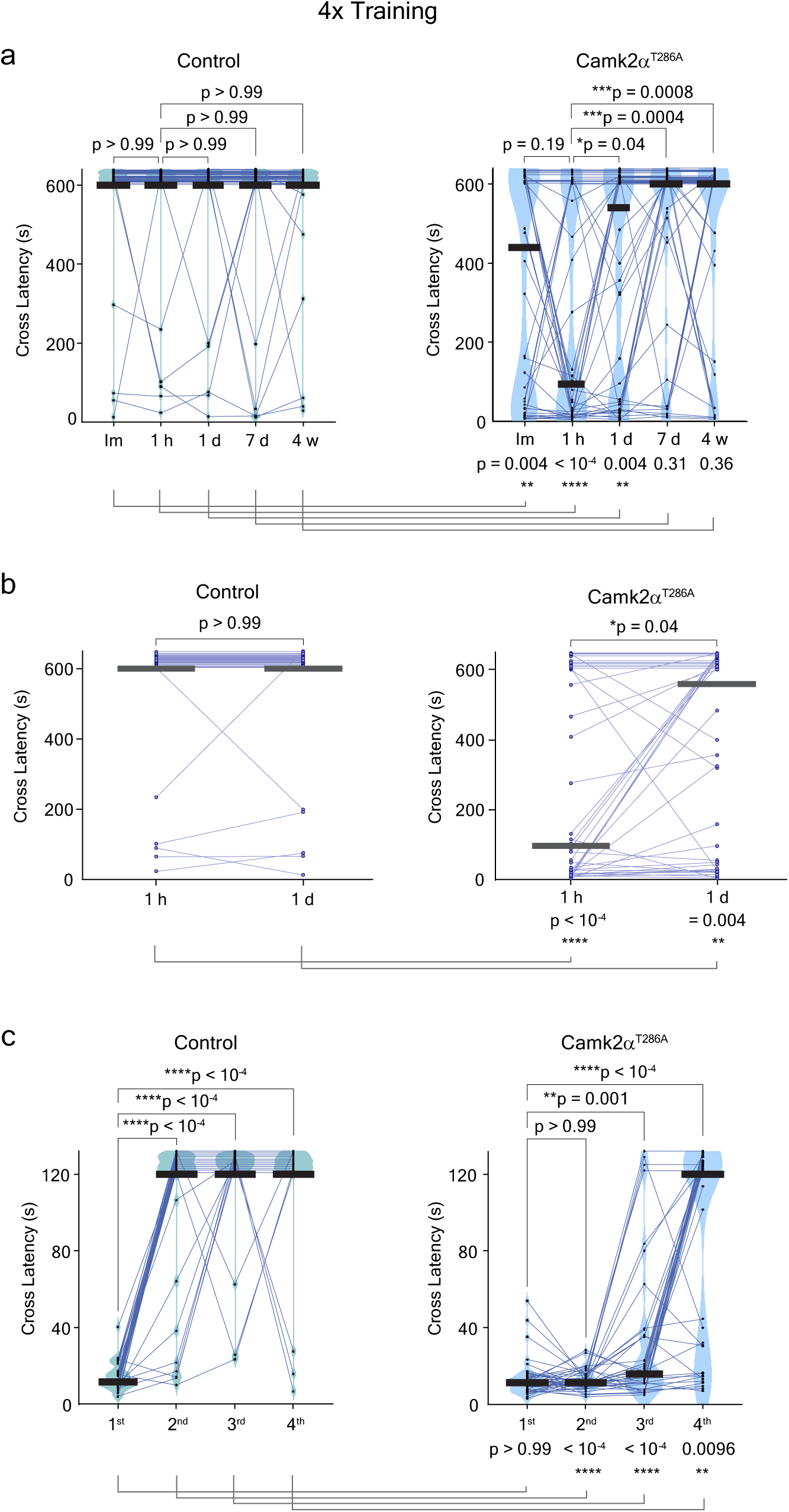
Performance of IA task for transgenic mice with inhibited CaMKII autophosphorylation with a maximum cross latency of 600 s. **a**, Performance of IA task immediately, 1 h, 1 d, 7 d and 4 w after 4 trials of training for *Camk2a*^T286A^ mice (right) or their littermate non-transgenic control (left). The experiment is similar to that in Fig. 2c, but with a longer maximum cross latency (600 s). **b**, Cross latencies at two selected time points (1 h and 1 d) from **a** are shown. **c**, Acquisition of IA memory for transgenic and control mice that were shown in **a-b**. Median is indicated by the horizontal bar. Two-sided Wilcoxon rank sum test was performed between 1 h and the other time points (**a-b**) or between the first and subsequent trials of training (**c**), respectively in each group. Mann-Whitney test was conducted between control and transgenic mice at each testing (**a-b**) or training sessions (**c**). n = 35 for control and 42 for *Camk2a*^T286A^ mice (**a-c**). *p < 0.05, **p < 0.01, ***p < 0.001, ****p < 0.0001 Data points with maximum latency values (600 s for **a-b**, 120 s for **c**) have been randomized between 601 and 640 (**a**) or 631 (**b**), or between 121 and 132 s (**c**).

**Supplementary Figure 6.**
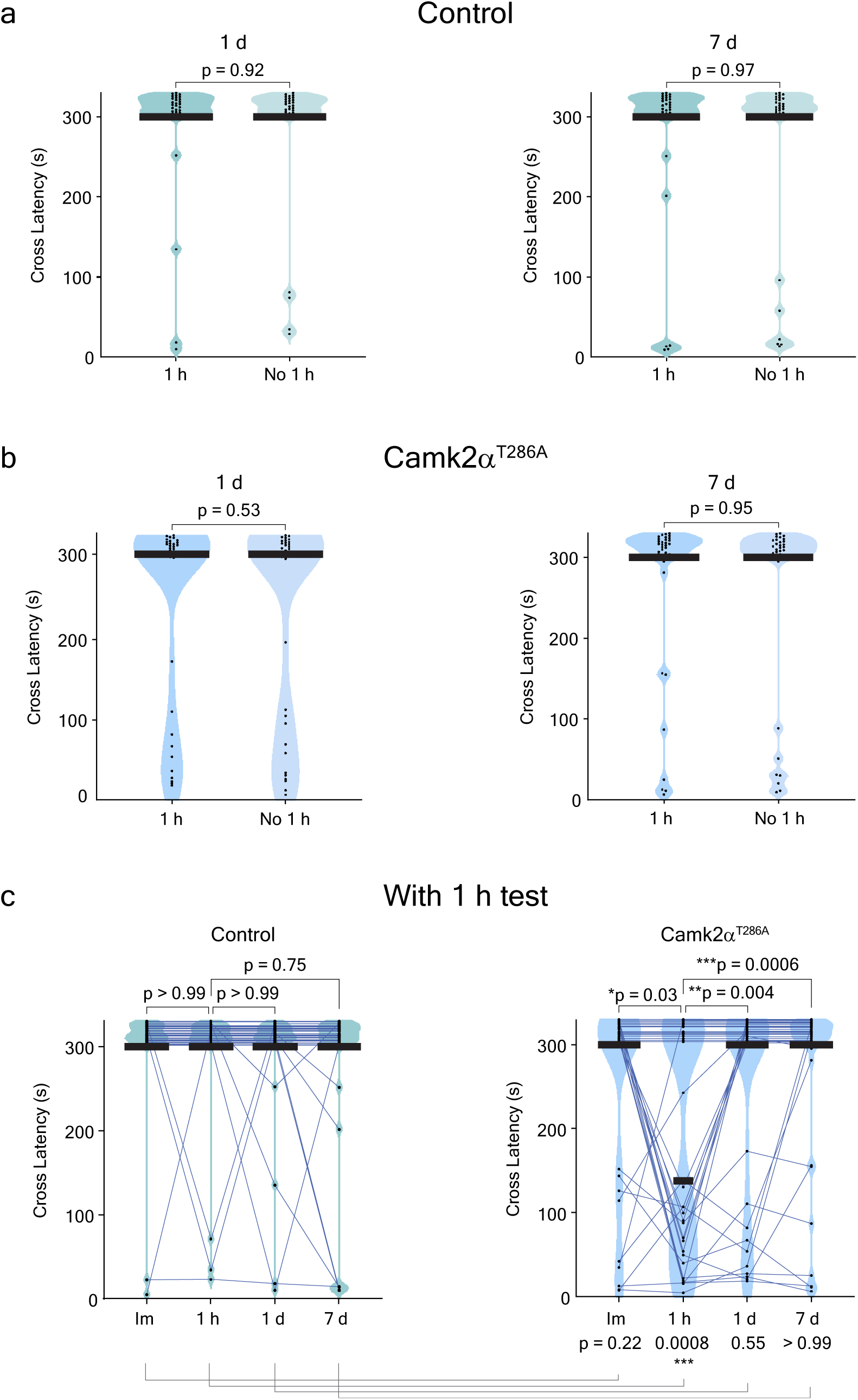
Performance of IA task for transgenic mice with inhibited CaMKII autophosphorylation in which the memory test 1 h after the IA training has been omitted. **a-b**, Performance of IA task 1 d (left) and 7 d (right) after 4 trials of training for *Camk2a*^T286A^ mice (**b**) or their littermate non-transgenic control (**a**). Each genotype was divided into 2 groups and for each group, a memory test was either conducted (‘1 h’ group) or omitted (‘No 1 h’ group) at 1 h from the training. **c**, Performance of IA task immediately, 1 h, 1 d, and 7 d after 4 trials of training for the ‘1 h’ group mice in both genotypes shown in **a-b**. Median is indicated by the horizontal bar. Mann-Whitney test was conducted between ‘1 h’ and ‘No 1 h’ groups (**a-b**) or between the ‘1 h’ control and transgenic mice at each time points (**c**). Two-sided Wilcoxon rank sum test was performed between 1 h and the other respective time points in each group (**c**). n = 31 for 1 h-control, 30 for No 1h-control, 35 for 1 h-*Camk2a*^T286A^, 33 for No 1h-*Camk2a*^T286A^ (**a-c**). *p < 0.05, **p < 0.01, ***p < 0.001 Data points with maximum latency values (300 s, **a-c**) have been randomized between 301 and 330.

**Supplementary Figure 7.**
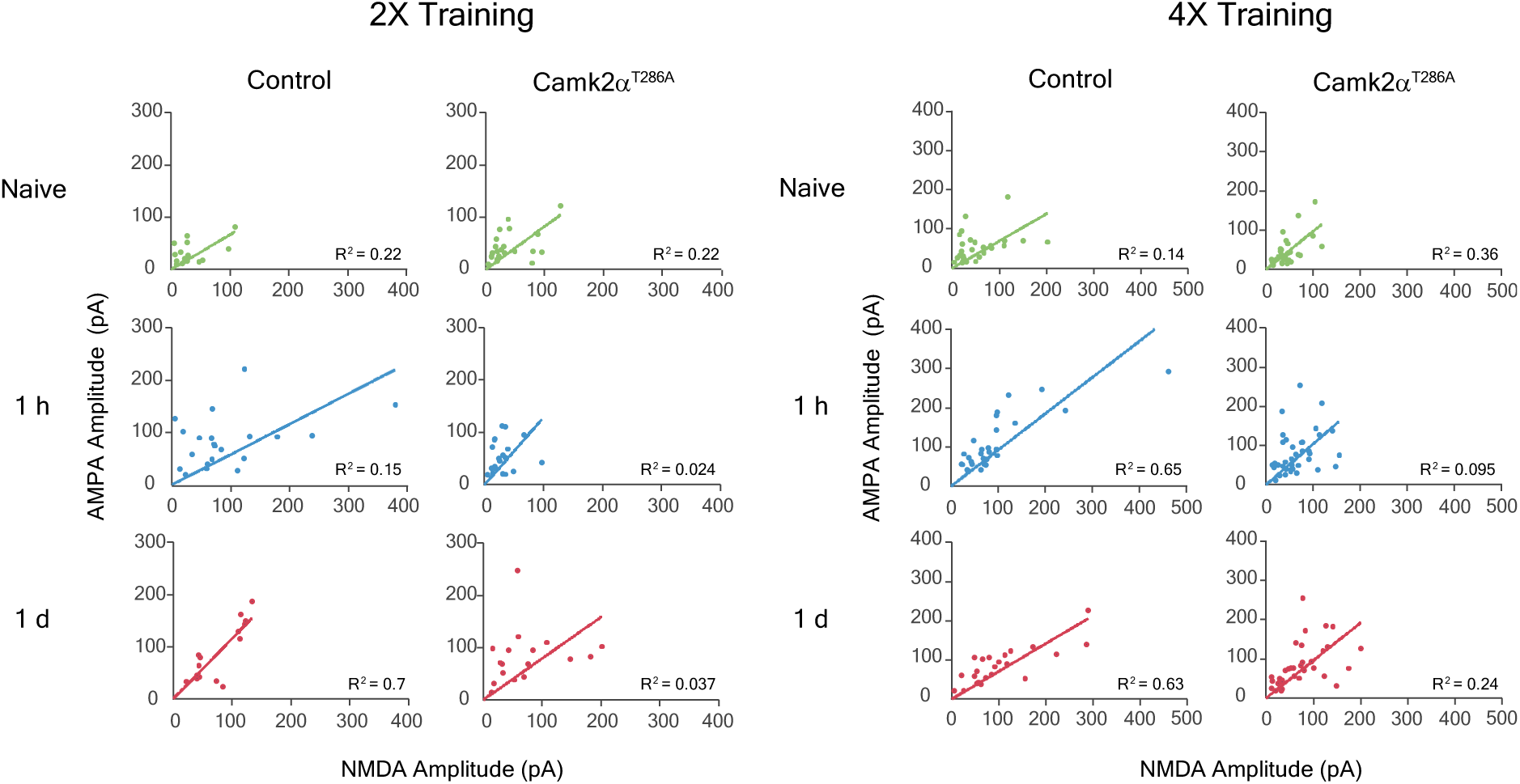
Correlation between AMPA and NMDA current amplitudes in acute brain slices obtained from transgenic mice with inhibited CaMKII autophosphorylation. Correlation between AMPA and NMDA current amplitudes in brain slices acquired from control and transgenic mice after 2 trial- and 4 trial-training IA. Values used in analysis were from the same set of cells shown in AMPA/NMDA ratio graphs in Fig. 3b and 3d. The square of the correlation coefficients (R^2^) and slopes from the nonlinear regression fit (solid line) are indicated. n_cells/animals_ for 2x training IA = 16/6 (control naïve), 21/5 (control 1 h), 15/5 (control 1 d), 22/11 (mutant naïve), 21/8 (mutant 1 h), 17/6 (mutant 1 d). n_cells/animals_ for 4x training IA = 29/6 (control naïve), 28/8 (control 1 h), 23/7 (control 1 d), 30/9 (mutant naïve), 37/11 (mutant 1 h), 34/13 (mutant 1 d).

**Supplementary Table 1.** Quantification of EGFP-P2A-paAIP2 expression in basolateral amygdala (BLA) complex in amygdala. Cells expressing the transgene within the BLA complex in both hemispheres were counted from a representative image for each mouse using ImageJ software. Percentage of the transgene expression as well as the number of the cells counted are shown for each animal. Cross latency values for all IA training and testing time points are shown for each mouse as well. n/a indicates that the transgene expression was not detectable by eyes under low magnification but slices were not available for later quantification.

| paAIP2neg<br>mouse ID | Transgene Expression |  |  |  | Cross Latency (s) |  |  |  |
| --- | --- | --- | --- | --- | --- | --- | --- | --- |
|  | Hemi 1 |  | Hemi 2 |  |  |  |  |  |
|  | % | GFP #/DAPI # | % | GFP #/DAPI # | 1<sup>st</sup> Training | 2<sup>nd</sup> Training | 1 h | 1 d |
| 1 | 37.4 | 450/1203 | 19.1 | 185/970 | 4.40 | 12.67 | 395.66 | 600.00 |
| 2 | 40.8 | 443/1087 | 0.0 | n/a | 8.54 | 14.81 | 146.21 | 439.17 |
| 3 | 53.2 | 604/1135 | 15.2 | 179/1174 | 10.14 | 120.00 | 600.00 | 600.00 |
| 4 | 33.7 | 366/1086 | 0.0 | 0/967 | 10.80 | 19.95 | 600.00 | 600.00 |
| 5 | 32.6 | 389/1192 | 33.6 | 398/1185 | 13.01 | 70.60 | 19.01 | 261.66 |
| 6 | 66.8 | 621/930 | 0.0 | 0/652 | 13.87 | 120.00 | 377.04 | 275.87 |
| 7 | 6.5 | 68/1053 | 14.6 | 125/854 | 15.81 | 120.00 | 600.00 | 600.00 |
| 8 | 34.3 | 523/1526 | 9.9 | 137/1377 | 16.54 | 34.16 | 600.00 | 600.00 |
| 9 | 13.9 | 126/905 | 0.0 | 0/675 | 16.88 | 49.38 | 600.00 | 600.00 |
| 10 | 22.4 | 242/1078 | 19.5 | 222/1141 | 18.01 | 37.10 | 600.00 | 317.38 |
| 11 | 87.1 | 412/473 | 0.0 | 0/1208 | 18.35 | 120.00 | 140.13 | 314.98 |
| 12 | 10.0 | 118/1180 | 2.4 | 34/1446 | 18.55 | 120.00 | 100.09 | 600.00 |
| 13 | 10.4 | 163/1573 | 86.2 | 1437/1667 | 20.88 | 116.51 | 600.00 | 600.00 |
| 14 | 34.9 | 372/1065 | 0.0 | 0/1295 | 21.08 | 120.00 | 600.00 | 600.00 |
| 15 | 35.7 | 524/1468 | 5.9 | 42/706 | 23.62 | 33.76 | 600.00 | 600.00 |
| 16 | 9.5 | 151/1596 | 6.2 | 58/936 | 26.69 | 120.00 | 600.00 | 600.00 |
| 17 | 10.2 | 117/1146 | 0.0 | 0/3237 | 30.42 | 120.00 | 103.83 | 600.00 |
| 18 | 41.9 | 427/1019 | 24.9 | 212/852 | 33.29 | 61.59 | 600.00 | 600.00 |
| 19 | 73.8 | 2140/2901 | 0.0 | 0/1720 | 33.76 | 120.00 | 600.00 | 600.00 |
| 20 | 37.4 | 207/553 | 64.5 | 395/612 | 35.03 | 120.00 | 600.00 | 146.94 |
| 21 | 20.2 | 399/1973 | 16.0 | 150/937 | 35.56 | 18.88 | 600.00 | 600.00 |
| 22 | 34.1 | 247/724 | 30.1 | 167/555 | 40.77 | 28.69 | 308.10 | 600.00 |
| 23 | 62.4 | 2237/3583 | 21.6 | 287/1326 | 43.24 | 120.00 | 600.00 | 600.00 |
| 24 | 73.7 | 837/1135 | 5.6 | 128/2291 | 44.64 | 120.00 | 321.11 | 600.00 |
| 25 | 22.6 | 317/1405 | 0.0 | 0/1436 | 45.31 | 58.52 | 600.00 | 58.19 |
| 26 | 12.3 | 85/693 | 4.8 | 24/502 | 45.37 | 28.36 | 600.00 | 378.64 |
| 27 | 20.9 | 227/1085 | 0.0 | 0/1302 | 48.78 | 120.00 | 600.00 | 91.82 |

**Supplementary Table 1.**
| paAIP2<br>mouse ID | Transgene Expression |  |  |  | Cross Latency (s) |  |  |  |
| --- | --- | --- | --- | --- | --- | --- | --- | --- |
|  | Hemi 1 |  | Hemi 2 |  |  |  |  |  |
|  | % | GFP #/DAPI # | % | GFP #/DAPI # | 1<sup>st</sup> Training | 2<sup>nd</sup> Training | 1 h | 1 d |
| 1 | 31.8 | 373/1172 | 20.8 | 254/1219 | 7.27 | 22.22 | 600.00 | 600.00 |
| 2 | 64.6 | 1318/2041 | 22.5 | 667/2958 | 8.07 | 120.00 | 600.00 | 600.00 |
| 3 | 54.4 | 271/498 | 0.0 | 0/577 | 8.27 | 22.28 | 178.37 | 600.00 |
| 4 | 49.6 | 348/701 | 0.0 | 0/487 | 8.67 | 120.00 | 76.74 | 600.00 |
| 5 | 37.5 | 200/533 | 7.7 | 47/612 | 9.27 | 29.42 | 21.35 | 600.00 |
| 6 | 4.2 | 109/2622 | 0.0 | 0/3426 | 9.54 | 24.42 | 27.62 | 600.00 |
| 7 | 79.0 | 722/914 | 7.0 | 29/412 | 11.47 | 13.34 | 18.48 | 600.00 |
| 8 | 42.3 | 341/807 | 17.8 | 54/304 | 11.74 | 120.00 | 167.96 | 600.00 |
| 9 | 18.5 | 94/509 | 0.0 | 0/1037 | 12.34 | 43.77 | 600.00 | 46.57 |
| 10 | 12.5 | 107/857 | 0.0 | 0/564 | 12.47 | 25.89 | 437.10 | 600.00 |
| 11 | 3.3 | 47/1428 | 52.0 | 1062/2042 | 12.81 | 120.00 | 55.65 | 600.00 |
| 12 | 20.8 | 877/4226 | 2.2 | 77/3438 | 14.01 | 120.00 | 600.00 | 600.00 |
| 13 | 40.4 | 392/971 | 0.0 | n/a | 14.14 | 120.00 | 401.53 | 600.00 |
| 14 | 21.0 | 271/1291 | 35.9 | 379/1055 | 15.94 | 20.61 | 72.67 | 512.44 |
| 15 | 3.2 | 86/2719 | 54.6 | 1835/3361 | 16.54 | 32.49 | 600.00 | 600.00 |
| 16 | 13.8 | 113/821 | 0.0 | 0/1027 | 16.68 | 120.00 | 42.17 | 600.00 |
| 17 | 58.1 | 241/415 | 55.3 | 347/627 | 18.08 | 78.87 | 600.00 | 600.00 |
| 18 | 42.2 | 231/548 | 48.9 | 595/1217 | 18.61 | 38.37 | 48.44 | 2.53 |
| 19 | 10.5 | 124/1186 | 78.8 | 469/595 | 18.81 | 20.68 | 91.08 | 349.54 |
| 20 | 17.5 | 179/1024 | 13.8 | 206/1493 | 19.48 | 16.81 | 26.55 | 600.00 |
| 21 | 44.0 | 521/1185 | 0.0 | n/a | 21.68 | 120.00 | 600.00 | 600.00 |
| 22 | 26.1 | 426/1634 | 42.9 | 1636/3816 | 27.22 | 120.00 | 25.42 | 600.00 |
| 23 | 24.8 | 206/831 | 28.9 | 301/1040 | 27.69 | 120.00 | 600.00 | 600.00 |
| 24 | 23.1 | 185/801 | 0.0 | n/a | 28.69 | 120.00 | 61.79 | 400.13 |
| 25 | 26.2 | 484/1845 | 11.2 | 59/528 | 29.42 | 120.00 | 66.73 | 600.00 |
| 26 | 40.3 | 898/2228 | 26.5 | 694/2622 | 29.96 | 120.00 | 600.00 | 270.53 |
| 27 | 11.2 | 115/1028 | 0.0 | 0/532 | 33.09 | 120.00 | 600.00 | 44.57 |
| 28 | 42.1 | 779/1849 | 7.5 | 32/428 | 33.56 | 120.00 | 148.01 | 600.00 |
| 29 | 71.4 | 2209/3092 | 0.0 | 0/2529 | 40.03 | 120.00 | 600.00 | 600.00 |
| 30 | 70.8 | 837/1182 | 0.0 | 0/829 | 40.90 | 120.00 | 256.18 | 600.00 |
| 31 | 42.0 | 591/1408 | 0.0 | 0/1402 | 42.30 | 42.57 | 81.88 | 157.42 |
| 32 | 20.9 | 159/761 | 26.0 | 953/3668 | 44.71 | 120.00 | 600.00 | 600.00 |
| 33 | 10.8 | 84/775 | 5.2 | 34/656 | 48.84 | 120.00 | 50.38 | 240.50 |

## References

1. Hebb, D, O. The Organization of Behavior: A Neuropsychological Theory. (Psychology Press, 1949).

2. Kandel, E. R., Dudai, Y. & Mayford, M. R. The molecular and systems biology of memory. Cell 157, 163–186 (2014).

3. Dudai, Y., Karni, A. & Born, J. The Consolidation and Transformation of Memory. Neuron 88, 20–32 (2015).

4. Hernandez, P. J. & Abel, T. The role of protein synthesis in memory consolidation: Progress amid decades of debate. Neurobiol. Learn. Mem. 89, 293–311 (2008).

5. Goto, A. et al. Stepwise synaptic plasticity events drive the early phase of memory consolidation. Science 374, 857–863 (2021).

6. Wally, M. E., Nomoto, M., Abdou, K., Murayama, E. & Inokuchi, K. A short-term memory trace persists for days in the mouse hippocampus. *Commun*. Biol. 2022 51 5, 1–11 (2022).

7. Izquierdo, L. A. et al. Molecular pharmacological dissection of short- and long-term memory. Cellular and Molecular Neurobiology 22, 269–287 (2002).

8. Sutton, M. A., Masters, S. E., Bagnall, M. W. & Carew, T. J. Molecular mechanisms underlying a unique intermediate phase of memory in Aplysia. Neuron 31, 143–154 (2001).

9. Gomis-González, M. et al. Protein Kinase C-Gamma Knockout Mice Show Impaired Hippocampal Short-Term Memory While Preserved Long-Term Memory. Mol. Neurobiol. 58, 617–630 (2021).

10. Silva, A. J., Paylor, R., Wehner, J. M. & Tonegawa, S. Impaired spatial learning in α-calcium-calmodulin kinase II mutant mice. Science 257, 206–211 (1992).

11. Giese, K. P., Fedorov, N. B., Filipkowski, R. K. & Silva, A. J. Autophosphorylation at Thr286 of the α calcium-calmodulin kinase II in LTP and learning. Science 279, 870–873 (1998).

12. Yamagata, Y. et al. Kinase-dead knock-in mouse reveals an essential role of kinase activity of Ca2+/calmodulin-dependent protein kinase IIα in dendritic spine enlargement, long-term potentiation, and learning. J. Neurosci. 29, 7607–7618 (2009).

13. Irvine, E. E., Vernon, J. & Giese, K. P. αCaMKII autophosphorylation contributes to rapid learning but is not necessary for memory. Nat. Neurosci. 8, 411–412 (2005).

14. Yamagata, Y., Yanagawa, Y. & Imoto, K. Differential involvement of kinase activity of Ca2+/ calmodulin-dependent protein kinase IIα in hippocampus-and amygdala-dependent memory revealed by kinase-dead knock-in mouse. eNeuro 5, (2018).

15. Murakoshi, H. et al. Kinetics of Endogenous CaMKII Required for Synaptic Plasticity Revealed by Optogenetic Kinase Inhibitor. Neuron 94, 37–47.e5 (2017).

16. Chang, J. Y. et al. CaMKII Autophosphorylation Is Necessary for Optimal Integration of Ca2+ Signals during LTP Induction, but Not Maintenance. Neuron 94, 800–808.e4 (2017).

17. Coultrap, S. J. et al. Autonomous CaMKII mediates both LTP and LTD using a mechanism for differential substrate site selection. Cell Rep. 6, 431–437 (2014).

